# Intracellular Aβ42 aggregation leads to cellular thermogenesis

**DOI:** 10.1101/2022.03.30.486355

**Authors:** Chyi Wei Chung, Amberley D. Stephens, Tasuku Konno, Edward Ward, Edward Avezov, Clemens F. Kaminski, Ali Hassanali, Gabriele S. Kaminski Schierle

## Abstract

The aggregation of Aβ42 is a hallmark of Alzheimer’s disease. It is still not known what the biochemical changes are inside a cell which will eventually lead to Aβ42 aggregation. Thermogenesis has been associated with cellular stress, the latter of which may promote aggregation. We perform intracellular thermometry measurements using fluorescent polymeric thermometers (FPTs) to show that Aβ42 aggregation in live cells leads to an increase in cell-averaged temperatures. This rise in temperature is mitigated upon treatment with an aggregation inhibitor of Aβ42 and is independent of mitochondrial damage that can otherwise lead to thermogenesis. With this, we present a diagnostic assay which could be used to screen small-molecule inhibitors to amyloid proteins in physiologically relevant settings. To interpret our experimental observations and motivate the development of future models, we perform classical molecular dynamics of model Aβ peptides to examine the factors that hinder thermal disspation. We observe that this is controlled by the presence of ions in its surrounding environment, the morphology of the amyloid peptides and the extent of its hydrogen-bonding interactions with water. We show that aggregation and heat retention by Aβ peptides are favoured under intracellular-mimicking ionic conditions, which could potentially promote thermogenesis. The latter will, in turn, trigger further nucleation events that accelerate disease progression.

## Introduction

Neurodegenerative diseases are a result of protein misfolding and aggregation, with ageing as its prime risk factor.^1^ This includes Alzheimer’s disease (AD), where the misfolding of amyloid-β (Aβ, in particular its 42-residue variant, Aβ42) into insoluble plaques is a characteristic of the disease. The most basic model for amyloid growth is a nucleation-elongation model, with nucleation being associated with a high energy barrier.^2^ There is a prevailing hypothesis of localised hotspots, i.e., energy-intensive areas, which may arise as the cell ages or during cellular stress. In these hotspots, the free energy activation barrier may be more likely overcome, hence spurring the initial nucleation, which subsequently facilitate monomeric addition (i.e., the elongation process). It has been shown, using *in vitro* isothermal titration calorimetric measurements, that Aβ42 elongation is an exothermic process^3,4^ which could lead to temperature gradients inside a cell. Mitochondrial thermogenesis, a heat-releasing effect linked to mitochondrial damage, the latter being associated with cell ageing and disease, may be another potential source of localised hotspots. It has long been established that Aβ42 interacts with, and can even localise and accumulate in mitochondria,^5,6^ thus resulting in mitochondrial dysfunction and neurotoxicity. Carbonyl cyanide p-(tri-fluromethoxy)phenyl-hydrazone (FCCP) is a proton uncoupler of oxidative phosphorylation which is commonly used to induce mitochondrial thermogenesis *in vitro*.^7,8^ It interferes the proton gradient across the electron transport chain (ETC), thereby disrupting adenosine triphosphate (ATP) synthesis. A recent study by Lautenschläger *et al*.^9^ has found that the addition of FCCP leads to greater intracellular Aβ42 aggregation in a HEK293T cell model. As mitochondria are the power source of the cell, it is unsurprising that oxidative stress incurred in the presence of Aβ42 aggregation could lead to changes in its metabolism, and hence to the overall energetics of the cell.

Hence, measuring intracellular temperatures is fundamental to a better understanding of the biochemical processes that may occur within a cell that is undergoing stress. Here, we perform intracellular thermometry using fluorescent polymeric thermometers (FPTs)^10^ and mitochondrial metabolism assays to investigate the intertwined pathways of Aβ42 pathology and mitochondrial dysfunction in a live cell model. The FPTs contain a fluorescence unit (*N*-{2-[(7-*N*,*N*-dimethylaminosulfonyl)-2,1,3-benzoxadiazol-4-yl](methyl)amino}ethyl-*N*-methylacrylamide; DBD-AA) which acts as a temperature sensor, as changes in its fluorescence lifetime are directly correlated to the changes in temperature.

We show here that the presence of Aβ42 in live HEK293T cells lead to an average temperature increase, in addition to inhibitions of glycolytic and oxidative respiratory pathways; however, the observed thermogenesis stems from Aβ42 elongation, instead of associated mitochondrial damage. This rise in temperature can therefore be mitigated upon treatment with a newly screened small molecule drug which binds to the C-terminus of Aβ, thereby causing steric hindrance and preventing its aggregation^11^. Lastly, using classical molecular dynamics simulations on Aβ, we show that heat dissipation from a protein is more greatly hindered in an intracellular-mimicking ionic environment compared to an aqueous one, due to altered protein-water hydrogen bonding interactions as a result of ionic interactions with termini groups on the protein. Furthermore, the three-dimensional packing of the Aβ in terms of the parallel and anti-parallel beta sheets also affects thermal relaxation behaviour of the protein.

## Results

### Intracellular thermometry can detect protein aggregation in live cells

In a first set of experiments, we incubated HEK293T cells with or without 500 nM unlabelled recombinant Aβ42^12^ for 24 hours prior to intracellular thermometry measurements. As additional controls, we treated cells with MJ040X, and Aβ42+ MJ040X, FCCP and Aβ42+FCCP. MJ040X is the the esterified version (i.e. to facilitate cellular uptake) of a recently discovered small molecule drug which sterically hinders the aggregation process by binding to the C terminus of Aβ42.^11^ Using time correlated single photon counting (TCSPC) fluorescence lifetime imaging microscopy (FLIM) to measure temperature-dependent changes in FPTs, we observe that the presence of Aβ42 elevates average intracellular temperatures by 2.8±0.6°C (**Figure 1**). Using super-resolution *direct* Optical Stochastic Reconstruction microscopy (*d*STORM), we confirm that exogenous addition of Aβ42 monomers and overnight incubation leads to the formation of fibrillar structures with an average length of 162 nm, dispersed throughout the cytoplasm of cells (**Supplementary Figure 1**). Furthermore, this thermogenesis effect is alleviated upon treatment with MJ040X to cells with exogenously added Aβ42, where temperature differences become insignificant from control cells which do not contain Aβ42. To confirm that MJ040X reduces the extent of Aβ42 aggregation in our current model, we used a fluorescence lifetime-based aggregation sensor, as we previously did to study amyloid aggregation in live cells ^13^. We added 10% HyLyte488 (HL488) labelled and 90% unlabelled Aβ42 to the cells prior to measuring the level of Aβ42 aggregation in the presence or absence of MJ040X. Upon addition of MJ040X, cells with added labelled Aβ42 have a higher fluorescence lifetime of ~0.1 ns in comparison to cells with Aβ42 without MJ040X, which indicates that Aβ42 is at a less aggregated state ^13^ in the presence of MJ040X (**Supplementary Figure 2**). Moreover, a non-cell based, *in vitro* assay to measure Aβ42 aggregation kinetics shows that MJ040 (i.e., the non-ester version of MJ040X and therefore the active MJ040 molecule present in the cytoplasm as the ester gets cleaved once the drug has reached the intracellular space^11^) works by inhibiting the exothermic elongation process rather than nucleation of Aβ42 (**Supplementary Figure 3**).

**Figure 1:**
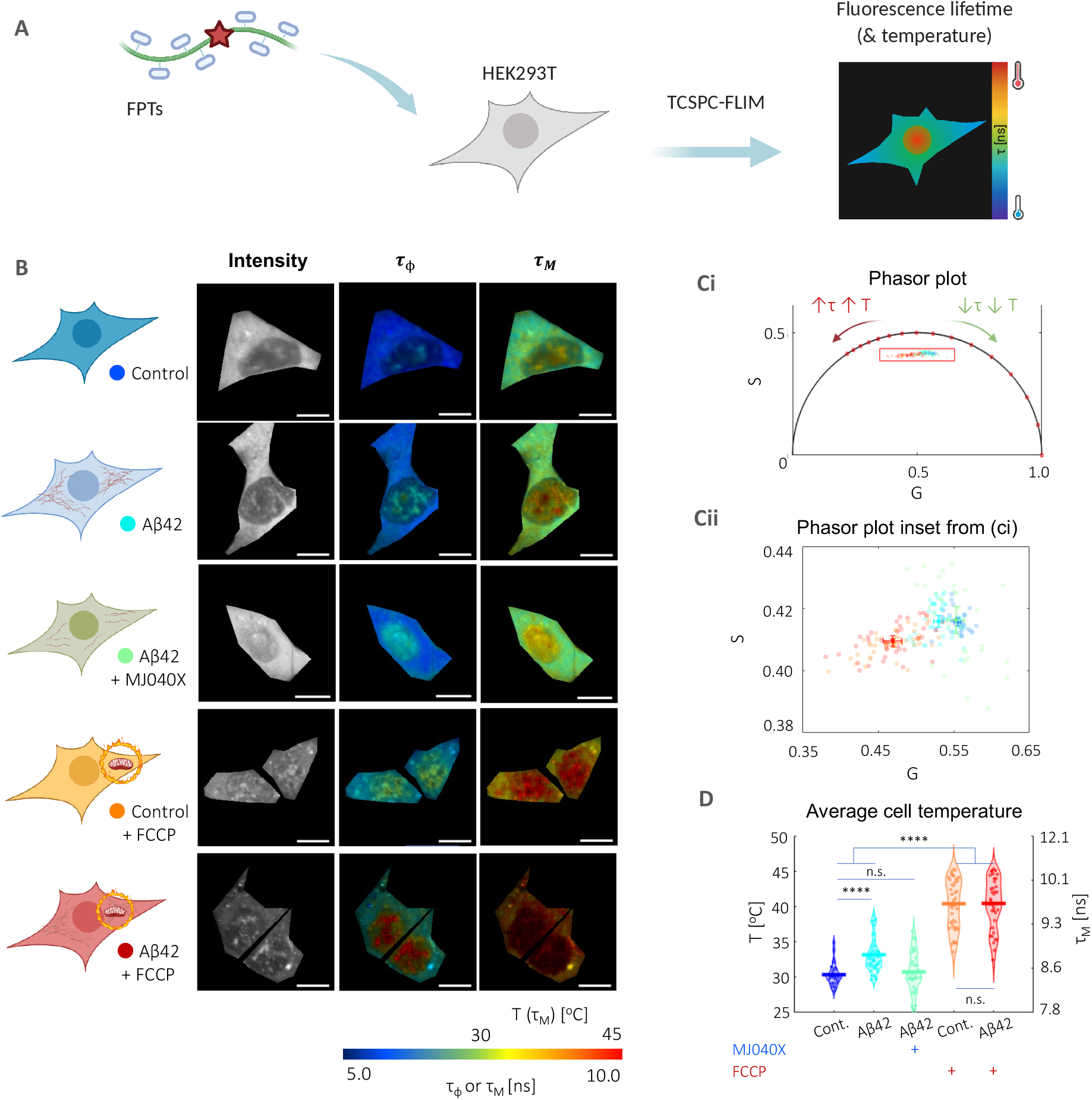
Intracellular thermometry using fluorescence lifetime-based readings from FPTs can be used as a platform for detecting Aβ42 aggregation and testing anti-aggregation drugs. (A) Cartoon representation of FPT-FLIM, where FPTs are introduced into live HEK293T for intracellular thermometry measurements based on fluorescence lifetime readouts. (B) Intracellular thermometry maps reveal the existence of intracellular temperature gradients. For each cell sample tested, their cartoon representation is shown alongside FPT-FLIM fluorescence intensity, and phase (τ_φ_) and modulation fluorescence lifetime (τ_M_) images, the latter being the temperature-calibrated parameter. (C) Biexponential decay of FPTs is apparent from significantly different τ_φ_ and τ_M_, as well as their phasors that fall within the universal semicircle (Ci), each dot and shaded circle represent the mean phasor from a single cell and standard deviation within the sample, respectively in the zoomed in plot (Cii). A legend is provided in the cartoon representations in (B). (D) Cell-averaged temperature values are given with mean and SEM average values of 30.4±0.5 (control), 33.2±0.8 (Aβ42), 30.8±0.9 (control+MJ040X), 40.4±1.2 (control+FCCP) and 40.4±1.2 °C (Aβ42+MJ040X). The presence of Aβ42 and/or FCCP elevated mean cell temperature slightly and significantly, respectively. The addition of MJ040X, which reduces the aggregation extent of Aβ42, reverts temperatures to that of the control. Thermometry experiments are conducted on >30 cells imaged over three biological repeats (i.e., N=3). One-way ANOVA tests (with Holm-Sidak’s multiple comparisons) were performed, where n.s. is not significant, ** is p<0.005, *** is p<0.001 and **** is p<0.0001. Cartoon representations are created on BioRender.com

For intracellular thermometry measurements, we validated our imaging setup using Rhodamine B, a dye with fluorescence emission properties that are strongly affected by temperature.^14,15^ (**Supplementary Figure 4A**). We assembled a temperature-controlled setup consisting of a stage top incubator and an objective warmer to control environmental parameters precisely. We then performed a temperature to fluorescence lifetime calibration of the FPTs in live cells (**Supplementary Figure 4B**). We analysed FPT-FLIM data using phasor plot, an *a priori* and global approach more suited to complex exponential decays (i.e., which the FPTs possess) and low photon count images.^16,17^ In comparison to conventional exponential fitting, which relies on non-linear least squares fitting, this involves the Fourier transform of time-domain TCSPC data into the frequencydomain, resulting in ‘phasors’ plotted on a polar plot. The bi-exponential decay of the FPTs^10^ is apparent from the difference in phase and modulation lifetimes (**Figure 1B&C**), and the latter is used as a calibration parameter to temperature due to its higher sensitivity to temperature (**Supplementary Figure 4B**). Cells with higher fluorescence lifetimes, hence higher temperatures, have phasors that move along the phasor plot in an anticlockwise manner (**Figure 1C&D**). The highest temperature differences (i.e., 10±1.0 °C) are measured in cells with added FCCP, a protonophore, showing the severe mitochondrial thermogenesis effect due to disruption in the ETC, as expected. There is no difference between FCCP-treated cells with and without Aβ42, indicating that exothermic effects from Aβ42 aggregation is outweighed by severe dysfunction of the mitochondria as triggered by the proton uncoupler (**Figure 1C&D**). Thus, FPT-FLIM can capture thermal events associated with Aβ42 aggregation in live cells.

### Exothermic elongation is the primary contributor to thermogenesis in cells with Aβ42 aggregation

By comparing the temperature elevation in the case of exogenously added Aβ42 cells, but with and without MJ040X, heating contributions can be attributed to exothermic elongation, as indicated above, but also by mitochondrial dysfunction. Hence to decouple these factors, we investigate mitochondrial metabolism using two different assays. The first one involves Ateam1.03, a CFP-mVenus Förster Resonance Energy Transfer (FRET) pair, which acts as a cytosolic adenosine triphosphate (ATP) sensor.^18^ The presence of ATP causes more interactions between the FRET pair, which can be detected by a decrease in the donor fluorescence lifetime. The results show that Aβ42 aggregation causes a significant loss of ATP which can be rescued by the addition of MJ040X, however, in the case of the control only cell, MJ040X does not significantly affect ATP levels. In the case of FCCP addition to control cells (i.e., the positive control), there is a significant reduction in ATP synthesis, with a corresponding rise in fluorescence lifetime of ~0.4 ns (**Figure 2A**). These results are further corroborated by a Seahorse Mito Stress assay, the gold standard technique which distinguishes between energy metabolism from contributing glycolytic and oxidative phosphorylation pathways, by the measurements of extracellular acidification rates (ECAR) and oxygen consumption rates (OCR), respectively (**Figure 2B**). When measured over time, cells with exogenously added Aβ42 with (cyan line) and without MJ040X treatment (green line) have lowered ECAR and OCR, in comparison to the control cells with and without Aβ42 (dark and light blue lines respectively) upon addition of modulators (e.g., oligomycin, FCCP and antimycin A/rotenone) which target different complexes forming part of the ETC (**Figure 2Bi**). Moreover, it is interesting to note that upon exposure to Aβ42 for 24 hours, the basal (i.e., initial) respiratory rates are already significantly impacted at two-fold the rates of control cells without Aβ42, i.e., from 203.7±29.7 to 78.7±14.2 pmol min^-1^ norm^-1^ (OCR) and from 51.8±.8.0 to 19.4±3.5 pH min^-1^ norm^-1^ (ECAR), where norm denotes normalisation to the cell count in the control (**Figure 2Bii**). This outweighs any effect MJ040X yields on mitochondrial metabolism. Hence, combining this with intracellular thermometry measurements, we conclude the main heating effect seen in cells with added Aβ42 derives from the exothermic effects of amyloid elongation rather than from mitochondrial damage, the latter of which is present but does not affect thermogenesis in our case.

**Figure 2:**
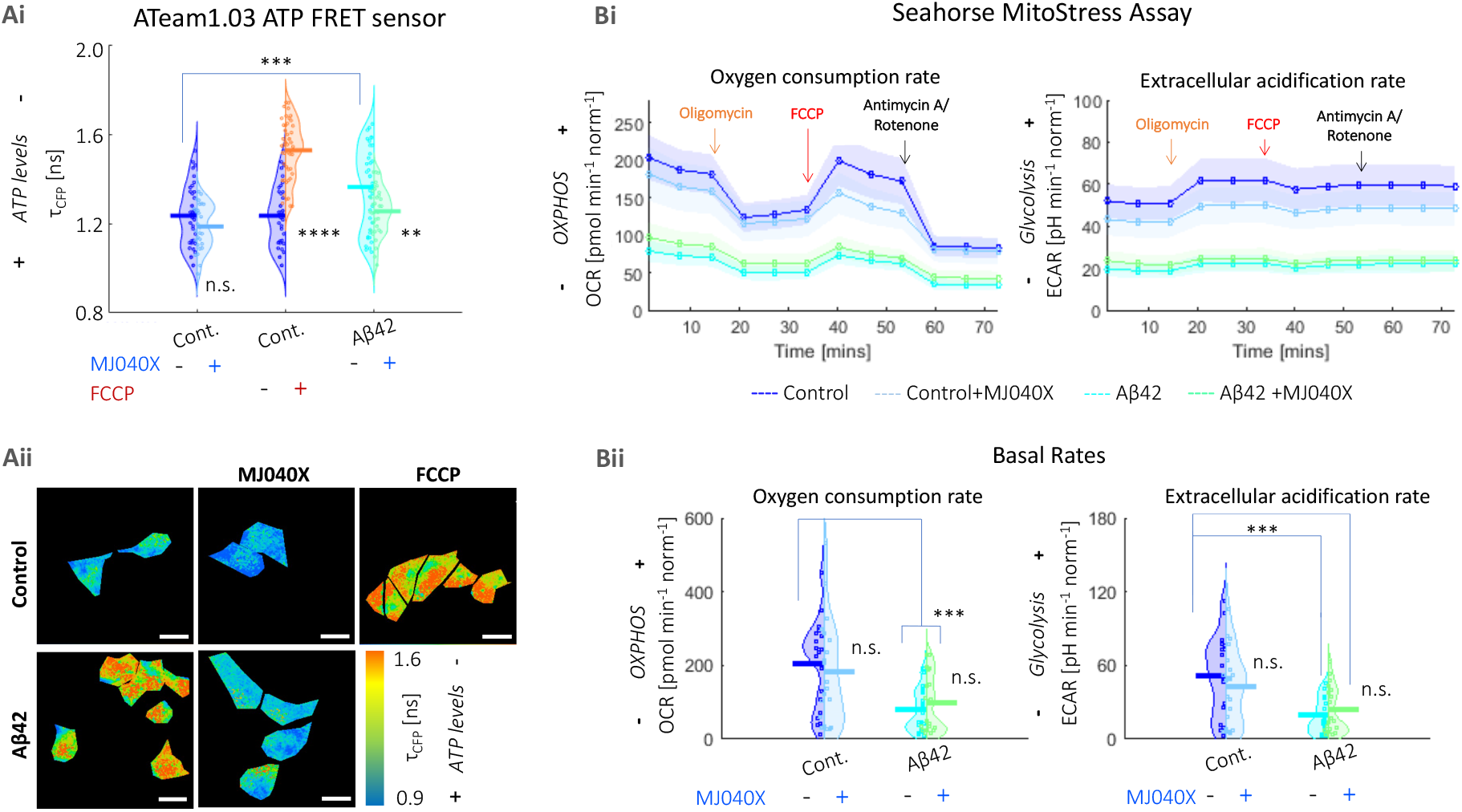
The presence of Aβ42 adversely impacts mitochondrial health. (Ai) Averaged fluorescence lifetimes and (Aii) corresponding maps of the fluorescence lifetime of the CFP donor (τ_CFP_) of the ATeam1.03 ATP sensor. ATP synthesis is negatively affected by the presence of Aβ42 and FCCP. The addition of MJ040X to cells with Aβ42 results in an increase in ATP production seen by the increase in τ_CFP_. (Bi) Seahorse assay shows lower OCR and ECAR (indicative of oxidative phosphorylation and glycolysis, respectively) over time, in the presence of Aβ42 (cyan for Aβ42 only and green for Aβ42+MJ040X), which prevails over the effect of MJ040X (blue for control+MJ040X and cyan for Aβ42+MJ040X) on mitochondrial metabolism. Arrows indicate when modulators are injected into cell media; this includes: oligomycin (orange), FCCP (red) and antimycin A/rotenone (black). (Bii) Basal OCR and ECAR (i.e., at the starting time point in bi) are impacted by the presence of Aβ42 after 24 hours of incubation. Basal OCR and ECAR are 203.7±29.7, 181.8±35.7, 78.7±14.2 & 79.1±19.4 pmol min^-1^ norm^-1^ and 51.8±.8.0, 43.1±7.5, 19.4±3.5 & 23.6±4.7 pH min^-1^ norm^-1^ respectively, for control, control+MJ040X, Aβ42 and Aβ42+MJ040X. Ateam1.03 analysis is based on >30 cells per sample over three biological repeats (N=3), and the Seahorse assay is based on the quantification of 4 wells per sample over 3 biological repeats (N=3) and are normalised by cell count. One-way ANOVA tests (with Holm-Sidak’s multiple comparisons) were performed, where n.s. is not significant, ** is p<0.005, *** is p<0.001 and **** is p<0.0001.

### Heat rentention by a protein is enhanced by ionic interactions and fewer hydrogen bonding interactions with water

We have thus far shown experimentally that non-equilibrium effects of Aβ42 elongation in a cell leads to an increase in its overall temperature. Our results suggest the importance of these effects in understanding the microscopic mechanisms associated with amyloid aggregation. Since it is currently impossible to model the non-equilibrium conditions in the cell, we turned to using empirical-based classical molecular dynamics to examine some factors (e.g., protein-water hydrogen bonding and ionic interactions) that may contribute to the fact that some of the heat released into the system may reside longer in certain areas of the cells which potentially leads to the observed thermogenesis effect. We thus investigated whether a) different amyloid structures or b) the local intracellular environment may favour the retention of heat in the system.

Aβ42 (like many misfolding proteins) possesses an intrinsically disordered nature, and the configuration which it adopts is influenced by ionic interactions with C- and N-termini and by wide networks of hydrogen bonding between protein and water.^19,20^ To simplify the case at hand, we first constructed molecular models based on different polymorphic segments of Aβ and investigate the effects of increased fibrillar size on heat disspation. We started from the base Aβ crystal structure of 2Y3J (Aβ30-35)^21^ and 2Y3L (Aβ38-42)^21^, which differ in the arrangement of their like-charged terminal groups, hence three-dimensional packing and arrangement of β-sheets: 2Y3J has a parallel β-sheet structure with like-charged terminal groups on the same side, whilst 2Y3L has an anti-parallel structure with the terminal groups alternating on each side. We concatenated them into separate 32 chain (32C) structures (**Figure 3Aii&B**). To represent Aβ at different stages of elongation, we further expanded our 2Y3J model to include single chain (1C, i.e., before the formation of β-sheets, **Figure 3Ai**) and 80 chain (80C, i.e., upon further elongation, **Figure 3Aii**) structures. They are then submerged in 4-site transferable intermolecular potential (TIP4P) water^22^ with and without the addition of 140 mM KCl, the latter as proxy to an intracellular environment. Under the application of an all-atom optimised potential for liquid simulations (OPLS/AA) force field,^23^ the system is energy minimised, followed by equilibration to a canonical ensemble (NVT) using two separate Nose-Hoover (N-H) thermostats^24,25^ on the protein and water at 300 K. Heating of the protein to 400 K is then performed using the N-H thermostat whilst the equivalent on water is kept at 300 K. The N-H thermostat of the protein is removed (whereas that of water is kept at 300 K) for production runs of thermal relaxation of the protein for 200 ps.

**Figure 3:**
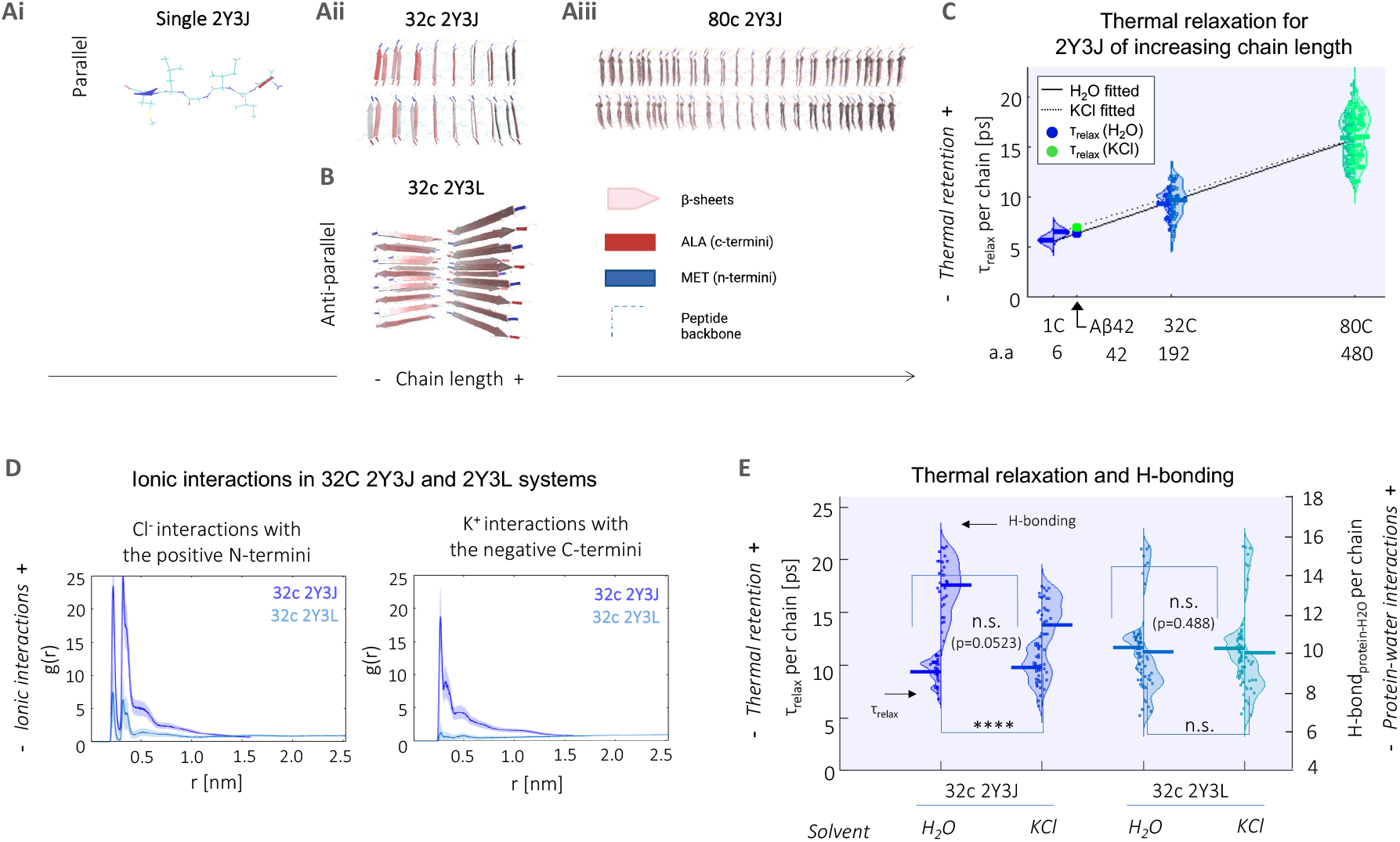
Heat retention by different Aβ structures, is favoured by fewer protein-water hydrogen bond interaction and in the presence of an ionic solvent. Structural illustration of model Aβ systems used, which include (A) parallel 2Y3J in (Ai) single, (Aii) 32C and (Aiii) 80C; as well as (B) anti-parallel 2Y3L. (C) Relaxation times (τ_relax_) at the single chain level increase with larger structures of 2Y3J, following a linear trend. For Aβ, predictions for τ_relax_ show heat retention is slightly enhanced in KCl (7.05 ps, green dot) compared to H_2_O (6.38 ps, blue dot). a.a. denotes number of amino acids.(D) Pair-correlation function (g(r)) shows that there is reduced ionic interaction with the charged terminal groups in anti-parallel 2Y3L, in comparison to parallel 2Y3J. (E) There is greater difference in both thermal retention (primary y-axis) and protein-water hydrogen bonding (secondary y-axis) between H_2_O and KCl in parallel 2Y3J than anti-parallel 2Y3L. t-test and one-way ANOVA tests (with Holm-Sidak’s multiple comparisons) are performed, where n.s. is not significant, ** is p<0.005, *** is p<0.001 and **** is p<0.0001.

Rajabpour *et al*.^26^ have compared modelling of solid-liquid interfaces, in their case, a silver nanoparticle-water system, using different molecular dynamics approaches and found that heat dissipation occurs at a time scale of ~5 ps and that conduction is the primary heat transfer mechanism in the nanoparticle surroundings at this time scale. As the Aβ crystal structures used are on the same length scale as a nanoparticle, we adopted a similar approach to study protein-water interfacial heat transfer and quantified relaxation time (τ_relax_, i.e., a measure of how quickly heat is dissipated from the protein, as calculated from exponentially decaying thermal relaxation curves) for each single chain comprising our model systems. We observe that single chain τ_relax_ increases with chain length of the concatenated 2Y3J system, i.e., from single to 32C and 80C (**Figure 3C**). We perform linear fitting for 2Y3J in both H_2_O (solid line) and KCl solvents (dotted line) and note that chain length seems to outweigh heat dissipation differences due to solvent effects especially at longer chain lengths in our model systems. For Aβ42 (which has 42 amino acids (a.a.)), we predict a single chain τ_relax_ of 6.38 ps in H_2_O (blue dot) and 7.05 ps in KCl (green dot), by interpolation of the linear fits. Alongside this, we also observe that there is slightly lower diffusivity of water molecules in the hydration shell in the 80C (0.33±0.01 in water and 0.32±0.02 x10^-5^ cm^-2^ s^-1^ in KCl solvent) compared to 32C structures (0.51±0.09 in water and 0.37±0.08 x10^-5^ cm^-2^ s^-1^ in KCl solvent), which is in itself already hindered when compared to the diffusivity of bulk water molecules (~2.8 x10^-5^ cm^-2^ s^-1^) (**Supplementary Figure 5**).

We next investigate the effects of ionic interactions both on heat dissipation and protein-water hydrogen bonding behaviour, using 32C structures of parallel 2Y3J and anti-parallel 2Y3L in KCl ionic solution. By computing cross-correlation functions (g(r)) of K^+^ and Cl^-^ ions with the negative C- and positive N-termini respectively, we observe that there are significantly lower ionic interactions in the case of 2Y3L, which is due to the weaker charge distribution across its structure. (**Figure 3D**). Consequently, the slight increase in τ_relax_ (p=0.0523) and decrease in protein-water hydrogen bonding for 2Y3J in KCl compared to in H_2_O, is not noticeable in the case of τ_relax_ (p=0.488) and hydrogen bonding for 2Y3L (**Figure 3E**). Hence, this demonstrates that heat dissipation from proteins is affected by ionic interactions, as well as hydrogen bonding interactions between protein and water. Moreover, by tracking the gyration radius of the molecules during the thermal relaxation process, the 32C 2Y3J takes on a more compact structure in KCl solution compared to water, giving a significant shrinkage in gyration radius of 0.2 nm (**Supplementary Figure 6**). This is analogous to experimental observation of proteins having more globular conformations (i.e., more aggregated structures) in the intracellular environment.^27^ However, this effect is less pronounced for 32C 2Y3L which we established has much weaker ionic interactions. Thus, thermal dissipation is hindered by increases in Aβ42 fibrillar structure and ionic interactions.

## Discussions

Aβ42 aggregation leads to thermogenesis which may impact cellular function. Although the presence of Aβ42 alone compromises mitochondrial health, we find that this does not result in mitochondrial thermogenesis sufficient to be detected by intracellular thermometry measurements, in contrast to when FCCP was used. We show, for the first time in live cells, that Aβ42 elongation is directly responsible for elevating cell-averaged temperatures. This could not only enhance the elongation of Aβ42 further, but could overcome the energy barrier for subsequent nucleation reactions, leading to the potential formation of oligomeric species which are more strongly associated with Aβ42-related pathology^28^. These newly formed structures could further enhance the thermogenesis effect as they act as seeds for elongation. Previous studies have revealed disease-related thermogenesis in cancerous cells^29^ and infected lesions^30^ using microcalorimetry measurements. On the other spectrum, it is interesting to note that hypothermia has been discussed as a potential neuroprotective therapy against dementia.^31,32^ Although the exact mechanism are still unknown, the overexpression of a cold-shock protein, RNA binding motif 3 (RBM3), results in reduced synaptic and neuronal loss in mouse models of neurodegeneration.

Moreover, the FPT-FLIM can also be used as a drug screening platform, as measured temperatures were sensitive to both our positive control (i.e., FCCP which triggers severe mitochondrial damage and thermogenesis) and negative control (i.e., MJ040 which inhibits Aβ42 aggregation). This method is particularly advantageous as it does not require any fluorescent labels to be attached to the amyloid protein which may hamper aggregation kinetics^33^ and/or affect the formation of different Aβ42 polymorphs^34^. The study further highlights the potential of small molecule drugs such MJ040. Since MJ040 is a potent inhibitor of Aβ42 elongation, it might be administered at later stages of the disease and thereby block further nucleation processes by reducing Aβ42 elongation-induced thermogenesis. This is strongly supported by *in vitro* studies which show that small temperature increases significantly enhance Aβ42 nucleation.^35,36^

One of the main detractions of reported intracellular thermometry measurements is the large temperature gradients, which over exceed predicted values from the heat-diffusion equation by a factor of 10^5^.^37^ One of the most controversial reports include Chretien *et al*.^38^ who, using MitoThermoYellow (a fluorescence-based mitochondrial-localised thermoprobe), observed that mitochondrial temperature is elevated up to 10 °C at its most active phase, without the use of any proton uncouplers. Nakano *et al*. ^7^ measured a 6—9 °C temperature increase across the whole cell by the addition of FCCP using a ratiometric thermoprobe (genetically encoded ratiometric fluorescent temperature indicator, gTEMP) transfected into HeLa cells; this agrees with the current work which has seen a ~10 °C increase in temperature with FCCP treatment in a different mammalian cell line, HEK293T. However, a similar study performed by Sugimura *et al*.^39^, by exploiting the temperature-sensitive OH stretching band of water using Raman spectroscopy, saw a more modest increase of 1.8 °C (and only in the cytosol) after FCCP treatment. Discrepancies in reported temperatures between different groups could arise due to variations in thermoprobe localisation, methods of performing the temperature calibration, experimental setup and methodology, and/or variations in cell cultures. In order to control for the latter, we performed the temperature calibration directly in the cells that were later used in the study, thus the sensor was already calibrated against potential cell to cell variability.

Semi-empirical models incorporating recent experimental findings made possible by advancement in the development of nano-/microscale thermoprobes, are useful for understanding the highly non-equilibrium environment inside a biological cell. In an attempt to understand our experimental findings, i.e. the Aβ42 elongation associated cellular thermogenesis, we investigated, using classical molecular dynamics simulation, how different Aβ42 structures and the ionic intracellular environment could promote heat retention. Proteins have thermal conductivities three to six times smaller than that of bulk water; in other words, they are able to sustain larger temperature gradients across their bodies.^40^ By creating model Aβ peptides of different chain length to mimic Aβ at different stages of elongation, we show that larger structures form fewer hydrogen bonds with water at a single chain level and heat dissipation occurs at a slower rate.

On the other hand, ionic interactions lead to amyloid proteins adopting a more globular configuration and are surrounded by hydration shells within which dynamic properties are distinct from bulk water. Here, retarded water mobility around large aggregates could thereby trigger the formation of potential hotspots. The hydrophobic effect also comes into play, as crowding effects effectively promote the overlap of hydration shells belonging to two different proteins. The latter has been show to result in the reduction to the overall hydration of the molecule and thereby triggers further aggregation.^27,41^ We further show that enhanced thermal retention in the presence of intracellular environment-mimicking KCl solution compared to water alone is due to ionic interactions and a decrease in protein-water hydrogen bonding interactions. However, further investigation (entailing non-equilibrium modelling) would be required to pinpoint the exact mechanisms which take place in an intracellular environment, where there is considerably higher packing density of proteins and other biomolecules.

## Conclusions

We show that Aβ42 elongation and not mitochondrial stress associated with Aβ42 aggregation is responsible for cellular thermogenesis as measured by FPTs, a highly sensitive temperature-dependent fluorescent sensor. We further validate that intracellular thermometry can be used as a tool to study Aβ42 aggregation and efficacy of an anti-aggregation small compound drug in a live cell model, thereby highlighting the potential of MJ040 as a therapeutic drug in AD.

We turn to classical molecular dyanmics to model different structures of Aβ, in an attempt to shed some light onto the mechanisms behind the non-equilibrium effects that give rise to heat dissipation from amyloids. We show that this is affected by the structural nature of the amyloid peptides, the ionic strength of its surrounding environment (highly prevalent to an intracellular environment), and the extent of hydrogen-bonding interactions of the amyloid with water. With the model above, we hope to spark off further work in this field to develop more accurate descriptors of complex intracellular phenomena that give rise to increased intracellular temperatures, which can be indicative of neurodegeneration.

## Methodology

### Cell culture

Human embryonic kidney cells (HEK-293T, American Type Culture Collection (ATCC), Manassas, VA, USA), cultured in a T75 cell flask with media, were incubated at 37 °C and 5% CO_2_. Media comprised of 90 v/v% Dulbecco’s modified Eagle’s medium (DMEM, ThermoFisher Scientific, Waltham, MA, USA), 10 v/v% foetal bovine serum (FBS, ThermoFisher Scientific), and 2 mM each of glutamax (ThermoFisher Scientific) and 2% penicillin-streptomycin (ThermoFisher Scientific). HEK-293T were passaged when 80-90% confluency was reached (i.e., twice a week). HEK-293T cells were plated into 8 well plates (IBIDI GmBH, Gräfelfing, Germany), and grown until 40—70% confluency before FPT introduction. For the cells with Aβ42, 500 nM solution of unlabelled A*β*42 monomer was added. For cells with MJ040X, 2.5 μM from a 100/200 μM stock in DMSO (Life Technologies, ThermoFisher Scientific) was added. Both Aβ42 and MJ040X were incubated for 24 hours before imaging. FCCP (Merck KGaA, Darmstadt, Germany) treatment was performed during the day of imaging, with FCCP added at a concentration of 10 μM (from a 10 mM stock in DMSO) and cells were left to incubate for 30 minutes. All reagents were kept at −20 °C and aliquoted to prevent freeze-thaw cycles.

### TCSPC-FLIM

Samples were imaged on a home-built confocal fluorescence microscope equipped with a time-correlated single photon counting (TCSPC) module. A pulsed, supercontinuum laser (Fianium Whitelase, NKT Photonics, Copenhagen, Denmark) provided excitation a repetition rate of 20 MHz (for FPT-FLIM) and 40 MHz (for imaging ATeam1.03 FRET sensor). This was passed into a commercial microscope frame (IX83, Olympus, Tokyo, Japan) through a 60x oil objective (PlanApo 60XOSC2, 1.4 NA, Olympus). The excitation and emission beams are filtered through select filters (Semrock, Rochester, NY, USA): (i) FF01-474/27-25 & FF01-515/LP-25 for FPTs and (ii) FF01-434/17 & FF01-470/28 for ATeam1.03. Laser scanning was performed using a galvanometric mirror system (Quadscanner, Aberrior, Gottingen, Germany). Emission photons were collected on a photon multiplier tube (PMT, PMC150, B&H GmBH, Berlin, Germany) and relayed to a time-corelated single photon counting card (SPC830, B&H GmBH). Images were acquired at 256×256 pixels for 200 s (i.e., 20 cycles of 10 s). Photon counts were kept below 1% of laser emission photon (i.e., SYNC) rates to prevent photon pile-up. TCSPC images were analysed using an in-house phasor plot analysis script (https://github.com/LAG-MNG-CambridgeUniversity/TCSPCPhasor), from which fluorescence lifetime maps and phasor plots were generated.

### FPT-FLIM

The linear cationic FPT (Funakoshi, Tokyo, Japan) was soaked overnight in Milli-Q water (Merck Millpore, Merck KGaA) to permit the full extension of the polymer, forming a 1 w/v% stock solution. On the day of imaging, the FPT stock solution was diluted in 5 w/v% glucose solution (made from a stock of 45% D-(+) glucose solution, Merck KGaA), to give a concentration of 0.03 w/v% FPT. Cell media was removed from a plated well and washed twice with 200 μL 1xPBS (ThermoFisher Scientific). 150 μL of 0.03 w/v% FPT solution was added to each cell well. After incubation at 25 °C without CO_2_ for 30 minutes, unincorporated FPT solution was removed, and each well was washed twice with 150 μL phosphate buffer solution (1×PBS). 250 μL of phenol red free Dulbecco’s Modified Eagle’s Medium (DMEM) (ThermoFisher Scientific), supplemented with 10% fetal bovine serum (FBS), was then added to the cell wells, ready for imaging.

Due to the relatively low fluorescence intensity of the FPT, a long-pass filter was used to maximise collection. The stage top heater (OKOLab, Ottaviano, Italy) was set at 30 °C, as the FPT are noted to clump at 37 °C; and at 5% CO_2_ and 20% relative humidity. An objective warmer (OW-1D, Multi-Channel Systems (MCS) GmbH, Reutlingen, Germany) was used to avoid the heat sink effect resulting from the significantly different thermal conductivities of the oil and bottom glass surface of the cell dish. This is controlled by a single channel single controller (TC-324C, MCS GmbH), which included a thermocouple (TA-29, MCS GmBH) that was inserted into the cell media through a small hole drilled on the lid to ensure cell media temperature is maintained at 30 °C. Results are based on >30 cells over three biological repeats.

Calibration of temperature to fluorescence lifetime was performed by temperature stepping between 28—45 °C (assumed temperatures were based on readings from the thermocouple inserted in the cell media), by adjusting temperature settings on both the OKOLab stage-top heater and objective warmer system. The final calibration equation based on modulation lifetime (*τ_M_*), in lieu of phase lifetime (*τ_φ_*) due to its higher sensitivity to temperature, as FPT phasors move more horizontally across the phasor plot (**Supplementary Figure 1b**).

We calibrated the FPTs in live HEK293T cells, subject to temperature stepping between 24—45 °C. The imaging setup was first validated using Rhodamine B (Merck KGaA), a standard fluorescence dye with fluorescence lifetimes that decrease as temperature increases.^15^ (see **Supplementary Figure 1**) For fluorescence lifetime, we applied phasor plot analysis *in lieu* of conventional exponential fitting, as the chosen is a non-statistical method that does not make *a priori* assumptions, hence permits more accurate complex exponential decay analyses. ^16,42^ Using calculated modulation lifetime based on phasors generated and the same definitions by Inada *et al*., ^10^ the system was found to have a temperature resolution of 0.8 °C, with modulation lifetimes ranging between 7.8 to 11.5 ns to cover a temperature range of 24 to 45 °C (see **Supplementary Table 1** for comparison).

### Ateam1.03 ATP FRET sensor

Design of the Förster resonance energy transfer (FRET-)based, cytosolic ATP sensor, Ateam1.03 can be found in Imamura *et al*.^43^. Two days before imaging, 200 ng Ateam1.03 plasmid diluted in antibiotics-free DMEM (ThermoFisher Scientific) was transfected using Lipofectamine 2000 (ThermoFisher Scientific) four hours after which the medium was changed, and the cells were kept incubated for another 20 hours before imaging. TCSPC-FLIM was only performed on the CFP donor channel whilst the YFP channel was used as a guide to confirm that the fluorescence seen in the CFP channel is due to Ateam1.03 rather than cell autofluorescence. ATeam1.03-nD/nA/pcDNA3 was a gift from Takeharu Nagai (Addgene plasmid number 51958; <u>RRID:Addgene_51958</u>). Analysis for the ATeam1.03 imaging, where SPCImage (B&H GmBH) is used to find the reduced fluorescence lifetime of CFP by bi-exponential fitting.

### Seahorse Mito Stress Assay

HEK293T were plated in 96 well plate (XF96, Agilent, Santa Clara, CA, USA), and 500 nM Aβ42 and/or 2.5 μM MJ040X were added alongside 24 hours before the Seahorse Assay. A Seahorse XFe96 analyser (Agilent) was used to perform the XF Cell Mito Stress Assay. Measurements had 18 mins intervals between drug addition, i.e., 1.5 μM oligomycin, followed by 1 μM FCCP and 0.5 μM rotenone/antimycin A (as part of Seahorse XF Cell Mito Stress Test Starter Pack, Agilent). Results are based on 3 biological repeats, with 5 wells plated per sample each time. Data were normalised to control according to cell count, as calculated by an automated cell counter (CountessII FL, ThermoFisher Scientific). Analysis was performed on Wave (Agilent, USA) before importing to MATLAB for plotting purposes.

### Molecular dynamics simulations

GROMACS (v2021, DOI: https://doi.org/10.5281/zenodo.4457626) was used for all simulations, with structures for Aβ30-35 (2Y3J) and Aβ35-42 (2Y3L) taken from the protein database. For concatenation of 2Y3J/L, Mercury (Cambridge Crystallographic Data Centre (CDCC), University of Cambridge) was used to alter the protein database (pdb) file. Each model is generated by fully solvating the Aβ peptide in TIP4P water within a triclinic box, with box edges at least 1 nm away from the peptide to satisfy periodic boundary conditions. For the KCl solvent, the equivalent of 140 mM of K^+^ and Cl^-^ ions were added, as all Aβ structures were of a neutral charge, the same number of cations and anions are added to preserve system neutrality. All systems were subject to an OLPS/AA forcefield and energy minimisation before being equilibration to a micromechanical (NVE) ensemble at 300 K for 100 ps. For disordered simulations, positional restraints were removed before energy minimisation. Heating of the peptide was achieved by introducing Nose-Hoover thermostat at 400 K and 300 K on the peptide and water, respectively, for 20 ps to generate initial conditions. Ordered and disordered simulations were performed to 200 ps and 10 ns, respectively, both at a timestep of 0.2 fs. Data output from simulations (in the form of .trr, .xvg) were then imported into MATLAB for batch analyses and producing figures and graphs. Relaxation time is calculated by linearising the exponentially decaying temperature profile (**Equation 1**). Hydration shells are assumed as water molecules at 0.3 nm away from the peptide.

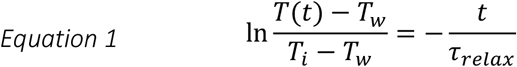

where ***T*** is temperature of the peptide at a given time, ***T_w_*** is temperature of the water (300 K), ***T_i_*** is temperature of the peptide at time zero (i.e., 400 K in most cases), ***t*** is time and ***τ_relax_*** is the thermal relaxation time. Thermal relaxation times and hydrogen bonding with water were calculated based on each chain within the peptide structure. Visualisation of the structures were produced using VMD.^44^

### Statistical analysis and plotting

All statistical analyses were performed on Prism 6 (GraphPad, San Diego, CA, USA), where a one-way ANOVA test with Holm-Sidak’s multiple comparison was applied. Violin plots were produced using open-source MATLAB code from Anne Urai (github.com/anne-urai).

## Data availability

Raw data is available through Cambridge Apollo Data Repository (DOI: 10.17863/CAM.82938)

## Code availability

TCSPC phasor plot analysis code is available via https://github.com/LAG-MNG-CambridgeUniversity/TCSPCPhasor. All other code is available upon request.

## Acknowledgements

This project received a pump priming grant from Alzheimer’s Research UK. C.W.C. is jointly funded by the Cambridge Trust and Wolfson College for her PhD. G.S.K.S. acknowledges funding from the Wellcome Trust (065807/Z/01/Z) (203249/Z/16/Z), the UK Medical Research Council (MRC) (MR/K02292X/1), Alzheimer Research UK (ARUK) (ARUK-PG013-14), Michael J Fox Foundation (16238) and Infinitus China Ltd.

## Contributions

G.S.K.S. and A.H. conceived the study. C.W.C. ran all experiments (unless otherwise stated), computational work and data analysis. A.D.S. purified Aβ42 protein and performed ThT assays and AFM. T.K. conducted the Seahorse assay and corresponding analysis. E.W. and C.W.C set up the TCSPC-FLIM system. E.A. provided feedback. C.F.K. helped with the design of the TCSPC-FLIM system and provided feedback. A.H. supervised on computational work. G.S.K.S. provided overall supervision and funding acquisition. C.W.C., A.D.S. and G.S.K.S. wrote and edited the manuscript.

## Additional information

Supplementary information (pdf)

## References

(1) Chiti, F.; Dobson, C. M. Protein Misfolding, Functional Amyloid, and Human Disease. Annu. Rev. Biochem. 2006, 75 (1), 333–366.

(2) Hoshino, M. Fibril Formation from the Amyloid-β Peptide Is Governed by a Dynamic Equilibrium Involving Association and Dissociation of the Monomer. Biophys. Rev. 2017, 9, 9–16.

(3) Kardos, J.; Yamamoto, K.; Hasegawa, K.; Naiki, H.; Goto, Y. Direct Measurement of the Thermodynamic Parameters of Amyloid Formation by Isothermal Titration Calorimetry. J. Biol. Chem. 2004, 279 (53), 55308–55314.

(4) Ikenoue, T.; Lee, Y.-H.; Kardos, J.; Yagi, H.; Ikegami, T.; Naiki, H.; Goto, Y. Heat of Supersaturation-Limited Amyloid Burst Directly Monitored by Isothermal Titration Calorimetry. Proc. Nat. Acad. Sci. USA 2014, 111 (18), 6654–6659.

(5) Hansson Petersen, C. A.; Alikhani, N.; Behbahani, H.; Wiehager, B.; Pavlov, P. F.; Alafuzoff, I.; Leinonen, V.; Ito, A.; Winblad, B.; Glaser, E.; Ankarcrona, M. The Amyloid β-Peptide Is Imported into Mitochondria via the TOM Import Machinery and Localized to Mitochondrial Cristae. Proc. Nat. Acad. Sci. USA 2008, 105 (35), 13145–13150.

(6) Devi, L.; Prabhu, B. M.; Galati, D. F.; Avadhani, N. G.; Anandatheerthavarada, H. K. Accumulation of Amyloid Precursor Protein in the Mitochondrial Import Channels of Human Alzheimer’s Disease Brain Is Associated with Mitochondrial Dysfunction. J. Neurosci. 2006, 26 (35), 9057–9068.

(7) Nakano, M.; Arai, Y.; Kotera, I.; Okabe, K.; Kamei, Y.; Nagai, T. Genetically Encoded Ratiometric Fluorescent Thermometer with Wide Range and Rapid Response. PLoS ONE 2017, 12 (2), e0172344.

(8) Hayashi, T.; Fukuda, N.; Uchiyama, S.; Inada, N. A Cell-Permeable Fluorescent Polymeric Thermometer for Intracellular Temperature Mapping in Mammalian Cell Lines. PLoS ONE 2015, 10 (2), e0117677.

(9) Lautenschläger, J.; Wagner-Valladolid, S.; Stephens, A. D.; Fernández-Villegas, A.; Hockings, C.; Mishra, A.; Manton, J. D.; Fantham, M. J.; Lu, M.; Rees, E. J.; Kaminski, C. F.; Kaminski Schierle, G. S. Intramitochondrial Proteostasis Is Directly Coupled to α-Synuclein and Amyloid β 1-42 Pathologies. J. Biol. Chem. 2020, 295 (30), 10138–10152.

(10) Inada, N.; Fukuda, N.; Hayashi, T.; Uchiyama, S. Temperature Imaging Using a Cationic Linear Fluorescent Polymeric Thermometer and Fluorescence Lifetime Imaging Microscopy. Nat. Protoc. 2019, 14, 1293–1321.

(11) Collins, S.; van Vliet, L.; Gielen, F.; Janeček, M.; Wagner Valladolid, S.; Poudel, C.; Fusco, G.; de Simone, A.; Michel, C.; Kaminski, C. F.; Spring, D. R.; Hollfelder, F.; Kaminski Schierle, G. S. A Unified in Vitro to in Vivo Fluorescence Lifetime Screening Platform Yields Amyloid β Aggregation Inhibitors. bioRxiv 2022.03.28.485913; doi: https://doi.org/10.1101/2022.03.28.485913 2021.

(12) Stephens, A. D.; Lu, M.; Fernandez-Villegas, A.; Kaminski Schierle, G. S. Fast Purification of Recombinant Monomeric Amyloid-β from E. Coli and Amyloid-β-MCherry Aggregates from Mammalian Cells. ACS Chem. Neurosci 2020, 11, 3213.

(13) Chen, W.; Young, L. J.; Lu, M.; Zaccone, A.; Strohl, F.; Yu, N.; Schierle, G. S. K.; Kaminski, C. F. Fluorescence Self-Quenching from Reporter Dyes Informs on the Structural Properties of Amyloid Clusters Formed in Vitro and in Cells. Nano Lett. 2017, 17 (1), 143–149.

(14) Kemnitz, K.; Yoshihara, K. Entropy-Driven Dimerisation of Xanthene Dyes in Non-Polar Solution and Temperature-Dependent Fluorescence Decay of Dimers. J. Phys. Chem. 1991, 95 (16), 6095–6104.

(15) Paviolo, C.; Clayton, A. H. A.; Mcarthur, S. L.; Stoddart, P. R. Temperature Measurement in the Microscopic Regime: A Comparison between Fluorescence Lifetime- and Intensity-Based Methods. J. Microsc. 2013, 250 (3), 179–188.

(16) Ranjit, S.; Malacrida, L.; Jameson, D. M.; Gratton, E. Fit-Free Analysis of Fluorescence Lifetime Imaging Data Using the Phasor Approach. Nat. Protoc. 2018, 13 (9), 1979–2004.

(17) Digman, M. A.; Caiolfa, V. R.; Zamai, M.; Gratton, E. The Phasor Approach to Fluorescence Lifetime Imaging Analysis. Biophys. J. 2008, 94 (2), L14.

(18) Imamura, H.; Nhat, K. P. H.; Togawa, H.; Saito, K.; Iino, R.; Kato-Yamada, Y.; Nagai, T.; Noji, H. Visualization of ATP Levels inside Single Living Cells with Fluorescence Resonance Energy Transfer-Based Genetically Encoded Indicators. Proc. Nat. Acad. Sci. USA 2009, 106 (37), 15651–15656.

(19) Jong, K.; Grisanti, L.; Hassanali, A. Hydrogen Bond Networks and Hydrophobic Effects in the Amyloid β 30-35 Chain in Water: A Molecular Dynamics Study. J. Chem. Inf. Model. 2017, 57 (7), 1548–1562.

(20) Bartolini, M.; Naldi, M.; Fiori, J.; Valle, F.; Biscarini, F.; Nicolau, D. v.; Andrisano, V. Kinetic Characterization of Amyloid-Beta 1-42 Aggregation with a Multimethodological Approach. Anal. Biochem. 2011, 414 (2), 215–225.

(21) Colletier, J. P.; Laganowsky, A.; Landau, M.; Zhao, M.; Soriaga, A. B.; Goldschmidt, L.; Flot, D.; Cascio, D.; Sawaya, M. R.; Eisenberg, D. Molecular Basis for Amyloid-β Polymorphism. Proc. Natl. Acad. Sci. USA. 2011, 108 (41), 16938–16943.

(22) Abascal, J. L. F.; Vega, C. A General Purpose Model for the Condensed Phases of Water: TIP4P/2005. J. Chem. Phys. 2005, 123 (23), 234505.

(23) Jorgensen, W. L.; Tirado-Rives, J. The OPLS Potential Functions for Proteins. Energy Minimizations for Crystals of Cyclic Peptides and Crambin. J. Am. Chem. Soc. 1988, 110 (6), 1657–1666.

(24) Nose, S. A Molecular Dynamics Method for Simulations in the Canonical Ensemble. Mol. Phys. 1983, 52 (2), 255–268.

(25) Hoover, W. G. Canonical Dynamics: Equilibrium Phase-Space Distributions. Phys. Rev. A 1985, 31 (3), 1695.

(26) Rajabpour, A.; Seif, R.; Arabha, S.; Heyhat, M. M.; Merabia, S.; Hassanali, A. Thermal Transport at a Nanoparticle-Water Interface: A Molecular Dynamics and Continuum Modeling Study. J. Chem. Phys. 2019, 150 (11), 114701.

(27) Stephens, A. D.; Kaminski Schierle, G. S. The Role of Water in Amyloid Aggregation Kinetics. Curr. Opin. Struc. Biol. 2019, 58, 115–123.

(28) Eckert, A.; Hauptmann, S.; Scherping, I.; Meinhardt, J.; Rhein, V.; Dröse, S.; Brandt, U.; Fändrich, M.; Müller, W. E.; Götz, J. Oligomeric and Fibrillar Species of β-Amyloid (Aβ42) Both Impair Mitochondrial Function in P301L Tau Transgenic Mice. J. Mol. Med. 2008, 86 (11), 1255–1267.

(29) Monti, M.; Brandt, L.; Ikomi-Kumm, J.; Olsson, H. Microcalorimetric Investigation of Cell Metabolism in Tumour Cells from Patients with Non-Hodgkin Lymphoma (NHL). Scand. J. Haematol. 1986, 36 (4), 353–357.

(30) Karnebogen, M.; Singer, D.; Kallerhoff, M.; Ringert, R. H. Microcalorimetric Investigations on Isolated Tumorous and Non-Tumorous Tissue Samples. Thermochim. Acta 1993, 229, 147–155.

(31) Peretti, D.; Bastide, A.; Radford, H.; Verity, N.; Molloy, C.; Martin, M. G.; Moreno, J. A.; Steinert, J. R.; Smith, T.; Dinsdale, D.; Willis, A. E.; Mallucci, G. R. RBM3 Mediates Structural Plasticity and Protective Effects of Cooling in Neurodegeneration. Nature 2015, 518 (7538), 236–239.

(32) Peretti, D.; Smith, H. L.; Verity, N.; Humoud, I.; de Weerd, L.; Swinden, D. P.; Hayes, J.; Mallucci, G. R. TrkB Signaling Regulates the Cold-Shock Protein RBM3-Mediated Neuroprotection. Life Sci. Alliance 2021, 4 (4), e202000884.

(33) Afitska, K.; Fucikova, A.; Shvadchak, V. v.; Yushchenko, D. A. Modification of C Terminus Provides New Insights into the Mechanism of α-Synuclein Aggregation. Biophys. J. 2017, 113 (10), 2182–2191.

(34) Wägele, J.; de Sio, S.; Voigt, B.; Balbach, J.; Ott, M. How Fluorescent Tags Modify Oligomer Size Distributions of the Alzheimer Peptide. Biophys. J. 2019, 116 (2), 227–238.

(35) Ghavami, M.; Rezaei, M.; Ejtehadi, R.; Lotfi, M.; Shokrgozar, M. A.; Abd Emamy, B.; Raush, J.; Mahmoudi, M. Physiological Temperature a Crucial Role in Amyloid in the Absence and Presence of Hydrophobic and Hydrophilic Nanoparticles. ACS Chem. Neurosci. 2013, 4 (3), 375.

(36) Sabaté, R.; Gallardo, M.; Estelrich, J. Temperature Dependence of the Nucleation Constant Rate in Beta Amyloid Fibrillogenesis. Int. J. Biol. Macromol. 2005, 35 (1-2), 9–13.

(37) Baffou, G.; Rigneault, H.; Marguet, D.; Jullien, L. A Critique of Methods for Temperature Imaging in Single Cells. Nat. Meth. 2014, 11 (9), 899–901.

(38) Chrétien, D.; Bénit, P.; Ha, H. H.; Keipert, S.; El-Khoury, R.; Chang, Y. T.; Jastroch, M.; Jacobs, H. T.; Rustin, P.; Rak, M. Mitochondria Are Physiologically Maintained at Close to 50 °C. PLoS Biol. 2018, 16 (1), e2003992.

(39) Sugimura, T.; Kajimoto, S.; Nakabayashi, T. Label-free Imaging of Intracellular Temperature by Using the O-H Stretching Raman Band of Water. Agnew. Chem., Int. Ed. Engl. 2020, 59, 7755–7760.

(40) Lervik, A.; Bresme, F.; Kjelstrup, S.; Bedeaux, D.; Miguel Rubi, J. Heat Transfer in Protein-Water Interfaces. Phys. Chem. Chem. Phys. 2010, 12 (7), 1610–1617.

(41) Chandler, D. Interfaces and the Driving Force of Hydrophobic Assembly. Nature 2005, 437 (7059), 640–647.

(42) Ranjit, S.; Malacrida, L.; Stakic, M.; Gratton, E. Determination of the Metabolic Index Using the Fluorescence Lifetime of Free and Bound Nicotinamide Adenine Dinucleotide Using the Phasor Approach. J. Biophotonics 2019, 12 (11), e201900156.

(43) Imamura, H.; Huynh Nhat, K. P.; Togawa, H.; Saito, K.; Iino, R.; Kato-Yamada, Y.; Nagai, T.; Noji, H. Visualization of ATP Levels inside Single Living Cells with Fluorescence Resonance Energy Transfer-Based Genetically Encoded Indicators. Proc. Natl. Acad. Sci. USA 2009, 106 (37), 15651–15656.

(44) Humphrey, W.; Dalke, A.; Schulten, K. VMD: Visual Molecular Dynamics. J. Mol. Graph. 1996, 14 (1), 33–38.

