## Supporting Information for "Intracellular Aβ42 aggregation leads to cellular thermogenesis"

### Supplementary Figures

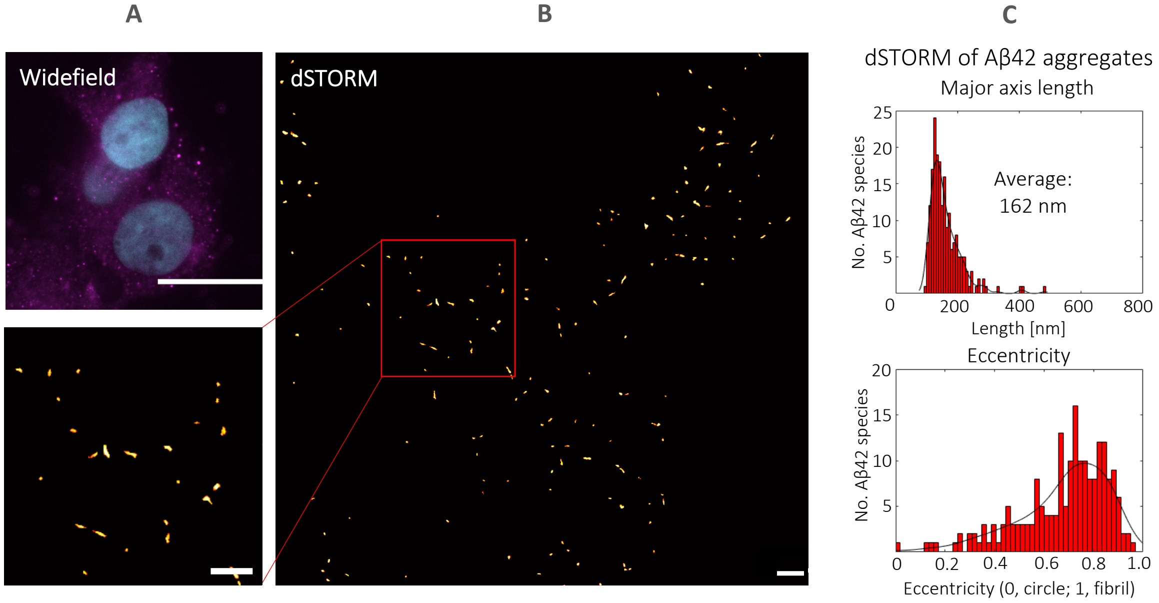

**Supplementary Figure 1: Exogenously added monomeric WT-Aβ42 forms fibrillar aggregates in HEK239T.** *(A) Widefield imaging provides insufficient resolution, as AF647 tagged antibody labelled Aβ42 (magenta) has dimensions below the diffraction limit. Cell nuclei are additionally stained with Hoechst 33342 to highlight Aβ42 formed only in the cytoplasmic regions. Scale bar, 10 µm. (B)* d*STORM images of AF647 tagged antibody labelled Aβ42 (red). Scale bars in main and zoomed image, 500 nm. (C) Morphological quantification in terms of major axis length (i.e., longest dimension length) and eccentricity (i.e., how fibrillar the structure is). 28 images were taken over 3 biological repeats.*

We incubate HEK293T cells with monomeric, unlabelled WT-Aβ42 for 24 hours to allow sufficient time for aggregate formation. The use of unlabelled protein provides better physiological relevance (i.e., avoids steric hindrance associated with fluorescent tags) and avoids any cross talk with the relatively dim FPT signal. As subsequent thermometry experiments are performed without being able to visualise Aβ42, a homogenous distribution of Aβ42 throughout cells was desired. Upon fixing the cells and immunostaining for Aβ42, we perform super-resolution *direct* Stochastic Optical Reconstruction Microscopy (*d*STORM). The technique works by localising fluorescence signal of a photoactivatable fluorophore (i.e., Alexa Fluor 647 in this case) undergoing photo-switching over 25000 frames captured at a high speed. Aβ42 aggregate structures formed appear primarily as elongated structures (i.e., with eccentricity values tending towards unity) with an average length of 162 nm, dispersed throughout the cytoplasmic region of the cells (**Supplementary Figure 1A**). Calculated dimensions agreed with values quoted by Esbjörner *et al.* [1], who visualised intracellular HiLyte-647 tagged Aβ42 in SH-SY5Y cells, using the same imaging setup. However, it should be noted that the aggregates formed were significantly smaller than the equivalent in the presence of pre-formed seeds. [2] Moreover, in comparison to the more aggressive familial mutant, E22G-Aβ42, structures formed do not display the same degree of polymorphism (i.e., ranging from clusters to bundles and large perinuclear aggresomes). [3]

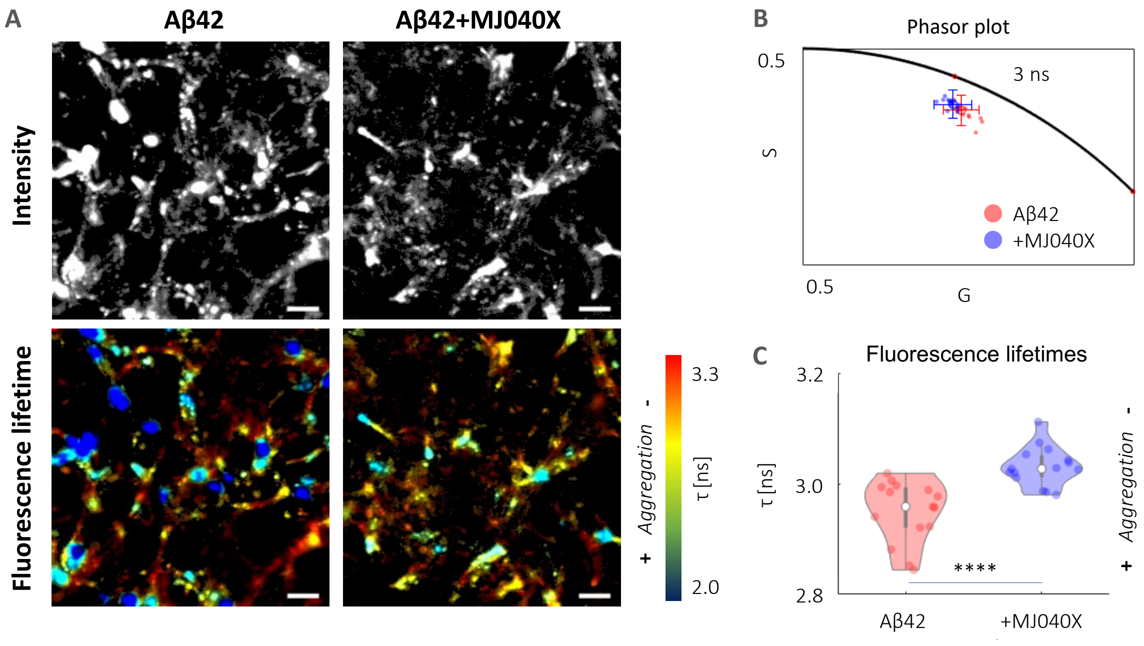

**Supplementary Figure 2: MJ040X, a small molecule drug, reduces the extent of Aβ42 aggregation.** *Experiments are based on measuring the fluorescence lifetime of 10% HyLyte488 tagged Aβ42 added to 90% unlabelled Aβ42. (A,B,C) Fluorescence lifetime maps, phasors, and averaged fluorescence lifetimes show that the addition of MJ040X alleviates Aβ42 aggregation. Based on 12 images collected over 3 biological repeats. Significance based on t-Test, where **** is p<0.0001.*

Ai

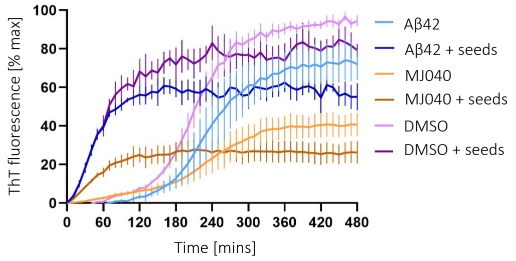

Aii

| | $t_{lag}$ [min] | $k$ [min$^{-1}$] |
|---|---|---|
| Aβ | 168.5 ± 20.5 | 0.52 ± 0.07 |
| MJ040 | 154.1 ± 30.4 | 0.25 ± 0.02** |
| DMSO | 138.6 ± 9.3 | 0.7 ± 0.02 |
| Aβ + seeds | | 0.72 ± 0.05 |
| MJ040 + seeds | | 0.29 ± 0.02*,* |
| DMSO + seeds | | 0.77 ± 0.06 |

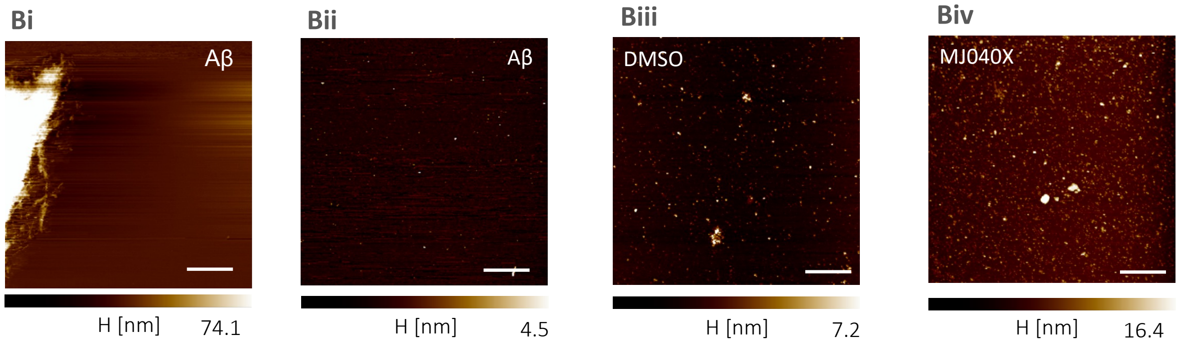

**Supplementary Figure 3: MJ040 significantly inhibits the elongation of Aβ42 in vitro.** *MJ040 was used in ThT-based assays as this is the ester cleaved form of MJ040X that would be found in cells inhibiting Aβ42 aggregation. ThT-based aggregation assays show 50 µM MJ040 inhibits elongation of Aβ42 in the presence and absence of 10% Aβ42 seeds. (Ai) 10 µM Aβ42 in 170 mM NaCl, 30 mM Tris, pH 7 with 20 µM ThT in a 368-well plate was incubated at 37°C for 8 hours. ThT fluorescence intensity was measured every 10 minutes, with double orbital agitation at 300 rpm for 15 seconds before each read. ThT fluorescence is presented as percentage maximum, averaged from three wells per plate, the experiment was repeated three times. The equivalent volume of DMSO (v/v) to MJ040 was added as a control. (Aii) The time to form fibrils ($t_{lag}$) and the elongation rate (k) were calculated from a linear fit of the exponential phase of the fibril growth curves. MJ040 significantly inhibited the elongation rate, (* is p<0.025 compared to Aβ42 + seeds and DMSO + seeds and ** is p<0.0073 compared to v/v DMSO using a one-way ANOVA with Dunnet's multiple comparison test). (B) Representative AFM images show the morphology of the samples after ThT-based assays. (Bi) The Aβ42 only sample showed presence of fibril clusters, (Bii) but also the presence of small fibrils and oligomers. For (Biii) DMSO and (Biv) MJ040 only oligomers were detected.*

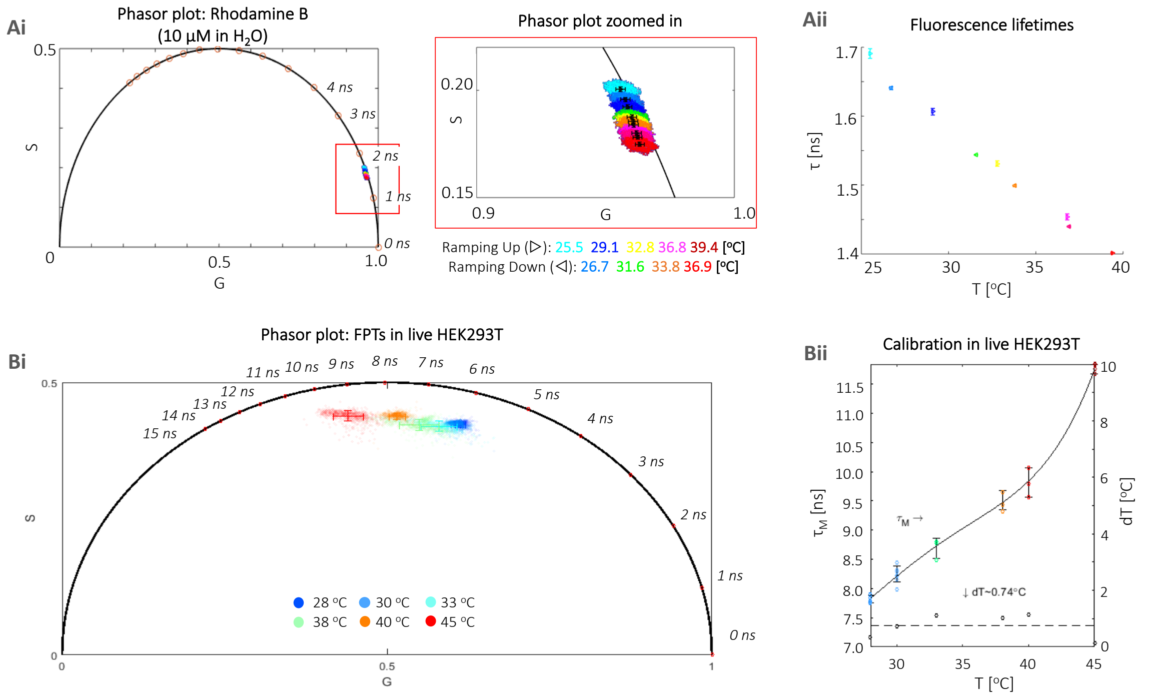

**Supplementary Figure 4: Temperature calibration of FTPs in live cells reveals a temperature resolution of 0.8 ºC.** *(A) The system was first validated with Rhodamine B, where a decrease of fluorescence lifetimes with increasing temperature was observed as seen in both the phasor plot (Ai) and corresponding fluorescence lifetime values (Aii). (b) FPT-FLIM calibration shows a horizontal, anticlockwise trajectory of FPT phasors with increasing temperatures (Bi), hence modulation lifetimes were used to calibrate the system to a temperature resolution of 0.74 ºC between 28—45 ºC (Bii).*

Temperature was determined by a thermocouple inserted into the cell medium, and an objective warmer (to prevent the heat sink effect by the objective) was used in conjunction with a stage top heater. To validate the ability of the system setup in terms of temperature stability and ramping, we first tested the system with Rhodamine B, a standard fluorescence dye that has a temperature-dependent fluorescence lifetime readout due to changes in the mobility of its diethylamino groups[4]. Measured fluorescence lifetime was reduced from 1.7 ns at 25 °C to 1.4 ns at 45 °C, agreeing with values in the literature.[5] Calibration in live cells is considered the more valid method, as it mirrors the actual experimental setup, however it is more time-consuming and difficult to perform. Hence, in addition, calibration in a cell extract solution (recommended as the easier method by Inada *et al.*[6]) was also attempted; however, clumping of the dyes in the cell extract became visibly apparent above 35 °C and fluorescence lifetime readings using TCSPC-FLIM became unreliably low in values. Hence, it is believed that cell extract calibration is only feasible in a spectrophotometer, a method more sensitive than TCSPC-FLIM and that does not involve inverted imaging, where FPT clumps become unavoidable.

*Supplementary Table 1: Comparison of FPT-FLIM data where fluorescence lifetimes are quoted as mean of bi-exponential lifetimes (exponential fitting) and modulation lifetimes (phasor plot)*

| | | Reported [6,7] | This work |
|---|---|---|---|
| Temperature range ($^{o}C$) | | 24—42 | 24—45 |
| Temperature resolution ($^{o}C$) | | 0.5 | 0.8 |
| Fluorescence lifetime [ns] | Exponential fitting* | 5—8.5 | 5—7.5 |
| | Phasor plots** | N/A | 7.8—11.5 |

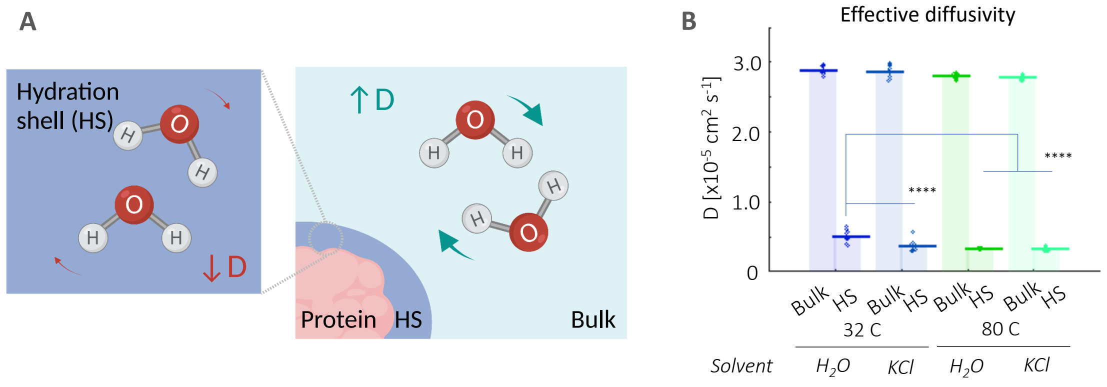

***Supplementary Figure 5: Large aggregates are more likely to promote heat retention.*** (A) Water molecules in HS that surround proteins submerged in solution have dynamic properties that differ from those in bulk water, e.g., lowered diffusivity in HS. Created on BioRender.com. (B) Effective diffusivity (D) values calculated from MSDs see up to a three-fold decrease in the HS compared to the bulk in both the 32 and 80C systems. There is a slight but significant effect of ionic KCl solution on hindering effective diffusivity in the 32 C system. Mean and standard deviation values of effective diffusivity are 2.89±0.05 (32C $H_2O$), 2.87±0.09 (32C KCl), 2.80±0.04 (80C $H_2O$), 2.78±0.03 x$10^{-5}$ cm$^{-2}$ s$^{-1}$ (80C KCl) for the bulk; and 0.51±0.09 (32C $H_2O$), 0.37±0.08 (32C KCl), 0.33±0.01 (80C $H_2O$), 0.32±0.02 x$10^{-5}$ cm$^{-2}$ s$^{-1}$ (80C KCl) for the HS. One-way ANOVA (Holm-Sidak's multiple comparison) is performed, where **** is $p<0.0001$.

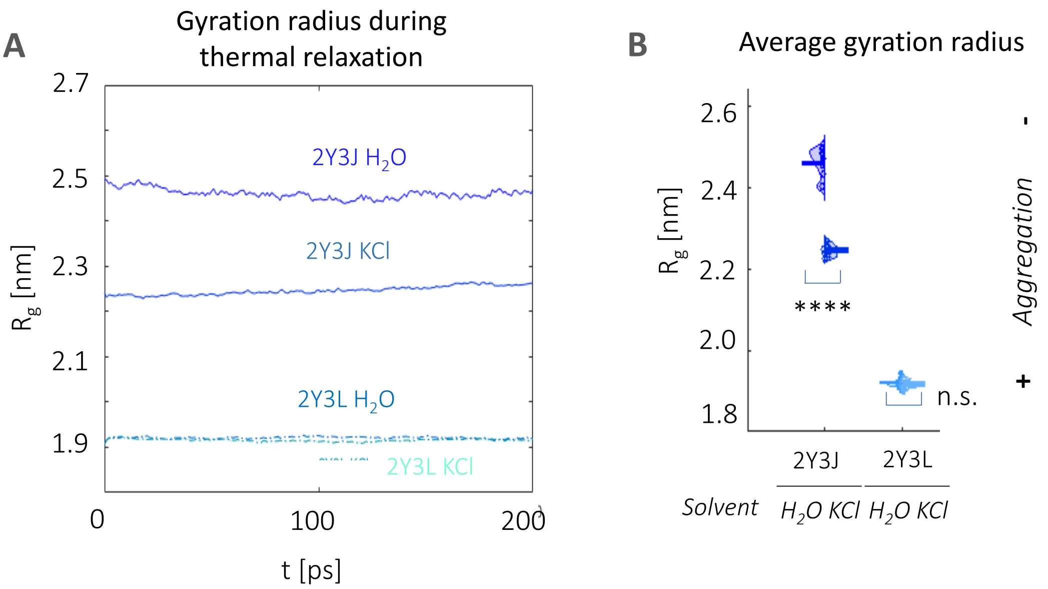

***Supplementary Figure 6: 32C 2Y3J is more aggregated in KCl than in water.*** (A) Gyration radius traces for 32C 2Y3J and 2Y3L in water and in KCl during the thermal relaxation process. (B) 2Y3J takes on a significantly, whilst 2Y3L a slightly more compact structure in KCl compared to water. T-test, where n.s. is not significant and **** is $p<0.05$.

### Supplementary methods

#### *AβM1-42 Purification*

Recombinant AβM1-42 (also referred to as Aβ) was purified as described in[8]. The plasmid pET3a containing AβM42 cDNA was transformed into *Escherichia coli* (*E. coli*) One Shot BL21 (DE3) pLysS (ThermoFisher Scientific). Liquid culture of *E. coli* was induced for expression for 4 hours when the $OD_{600}$ reached 0.6–0.8 by the addition of 1 mM isopropyl-β-thiogalactopyranoside (IPTG). The cells were pelleted in 50 mL volumes and the supernatant discarded. The pellet contained Aβ in inclusion bodies. The inclusion bodies were washed four times in wash buffer, as detailed in [8] to obtain clean and pure inclusion bodies. The inclusion body pellet from 50 mL of culture was placed on ice with a small magnetic stir bar on a magnetic stirrer. 200 µL of 6 M guanidinium chloride (GuHCl) was added to the pellet and stirred vigorously for 30 min to solubilise the Aβ. 15 mL of ice-cold ion exchange (IEX) buffer A (10 mM Tris, 1 mM ethylenediaminetetraacetic acid (EDTA), pH 9) was added slowly to the solubilised pellet to dilute the 6 M GuHCl and to permit binding of Aβ to the ion exchange column. The solubilised Aβ was filtered through a 0.22 µm filter (Millex-GP, MillPore, Merck KGaA) before being placed on ice prior to chromatography. Aβ was loaded onto a HiTrap Q HP column (GE, Healthcare, Chicago, IL, USA) and eluted against a linear gradient of IEX buffer B (10 mM Tris, 1 mM EDTA, 0.75 M NaCl, pH 9) over seven column volumes followed by two column volumes of 100% buffer B. Purification was performed on an ÄKTA Pure Fast Protein Liquid Chromatography (FPLC) and monitored by absorption at 280 nm (GE Healthcare). The concentration of Aβ was determined by absorption at 280 nm on a NanoVue spectrometer (Biochrom Ltd., Cambridge, UK) using the extinction coefficient of 1490 $M^{-1}$ $cm^{-1}$.

#### *Fixing of cells for the immunostaining of Aβ42*

Cell media was removed and replaced with 4% paraformaldehyde (PFA, Merck KGaA) diluted in 1×PBS. The sample was fixed for 10 minutes, before blocking with 5 w/v% bovine serum albumin (BSA, Merck KGaA) diluted in PBS for 1 hour. Between antibody incubation, three washes of 50 µM Triton X-100

(Invitrogen, ThermoFisher Scientific) in PBS (henceforth referred to as PBST) were performed. Primary and secondary antibodies used were β-Amyloid(1-42) polyclonal antibody (Invitrogen, ThermoFisher Scientific) and Goat anti-Rabbit Alexa Fluor 647 (ThermoFisher Scientific), diluted by 1:100 and 1:400 in PBST respectively. The sample was kept at room temperature throughout all the steps and wrapped in aluminium foil to prevent any bleaching especially after the addition of the secondary antibody. For storage before imaging, the sample was kept at 4 $^{o}$C in dark conditions.

#### d*STORM*

Before imaging the fixed cells, PBS is replaced with *d*STORM photo-switching buffer consisting of 50 mM Tris pH8 solution supplemented with 10 mM sodium chloride (ThermoFisher Scientific), 10% glucose (ThermoFisher Scientific), 50 mM monoethanolamine (MEA, Merck KGaA), 0.5 mg/mL glucose oxidase (Merck KGaA) and 40 µg/mL catalase (Merck KGaA). They are mounted on a custom-build microscope with an IX-73 Olympus frame (Olympus) with a 647 nm laser (VFL-P-300-647-OEM1-B1, MPB Communications Inc., Quebec, Canada). Laser light entering the microscope frame was reflected from a dichroic mirror (ZT647rpc, Chroma, Bellows Fall, VT, USA) onto a 100× 1.49 NA oil total internal reflection (TIRF) objective lens (UAPON100XOTIRF, Olympus), before reaching the sample. Light emitted by the sample passed through the dichroic and a set of 25 mm band-pass filters (FF01-680/42-25, Semrock) before reaching the microscope side port. Images were then relayed onto a camera (Andor iXon Ultra 897, Oxford Instruments, Belfast, UK) by a 1.3x magnification Twincam image (Cairn, Kent, UK). The image pixel size was measured to be 117 nm using a ruled slide. Each 256x256 image was acquired as stacks of 15000 images with an exposure time of 0.01 ms. Around eight images were captured per biological repeat. Fluorophore localisations are first detected using the Fiji plugin, ThunderSTORM[9], before reconstruction using an in-house MATLAB script detailed in the methodology section of [1]. Reconstructed images were then further analysed in a separate MATLAB script to quantify the major axis length (i.e., longest dimension) and eccentricity of Aβ42 aggregates. Aggregate masking was achieved by an intensity threshold, following a pixel-size threshold to ensure that only aggregates with dimensions above the resolution limit (i.e., estimated at 70 nm) were included in the analysis.

#### *ThT-based aggregation assays of recombinant Aβ42*

20 µM of freshly made thioflavin-T (ThT) (abcam, Cambridge, UK) was added to 10 µM Aβ in 170 mM NaCl, 30 mM Tris, pH 7. 50 µM of MJ040X or the equivalent volume of DMSO, which MJ040X is dissolved in, were added. 10% seeds were made by incubating 10 µM of Aβ for 24 hours and sonicating them for 5 seconds at 30% amplitude (Digital Sonifier® SLPe, model 4C15, Branson, Danbury, MA, USA). 25 µL of each sample was added in triplicate to a 368-well plate black plate with a clear bottom (Grenier Bio-One GmbH). The plates were sealed with a SILVERseal aluminum microplate sealer (Grenier Bio-One GmbH). Fluorescence measurements were taken using a FLUOstar Omega plate reader (BMG LABTECH GmbH, Ortenbery, Germany). The plates were incubated at 37 °C with double orbital shaking at 300 rpm for 15 seconds before each read every 10 minutes for 8 hours. Excitation was set at 440 nm, and the ThT fluorescence intensity was measured at 480 nm emission with a 2200 gain setting. ThT-based assays were repeated three times and the data normalised to the maximum fluorescence per plate. A linear trend line was fitted along the exponential phase of the ThT fluorescence curve and **Equation 1** was used to calculate the time to form the first fibrillary structures, lag time ($t_{lag}$), at the intercept of the x axis, and the elongation rate from the slope of the exponential phase ($k$), indicating growth rate.

*Equation 1*

$$y = kx - t_{lag}$$

#### *Atomic force microscopy on Aβ42*

The contents of wells from the ThT-based assays were deposited on a freshly cleaved mica surface for 20 min. The mica was washed three times in 18.2 MΩ.cm $dH_2O$ to remove lose protein. Images were acquired in $dH_2O$ using tapping mode on a BioScope Resolve (Bruker GmbH, Karlsruhe, Germany) using 'ScanAsyst-Fluid+' probes. 256 lines were acquired at a scan rate of 0.966 Hz per image with a field of view of 4 µm. Images were adjusted for contrast and exported from NanoScope Analysis 8.2 software (Bruker GmbH).

### References

1. Esbjörner, E. K. *et al.* Direct observations of amyloid β Self-assembly in live cells provide insights into differences in the kinetics of Aβ(1-40) and Aβ(1-42) aggregation. *Chem. Biol.* **21**, 732–742 (2014).

2. Kaminski Schierle, G. S. *et al.* In situ measurements of the formation and morphology of intracellular β-amyloid fibrils by super-resolution fluorescence imaging. *J. Am. Chem. Soc.* **133**, 12902–12905 (2011).

3. Lu, M. *et al.* Structural progression of amyloid-β Arctic mutant aggregation in cells revealed by multiparametric imaging. *J. Biol. Chem.* **294**, 1478–1487 (2019).

4. Kemnitz, K. & Yoshihara, K. Entropy-driven dimerisation of xanthene dyes in non-polar solution and temperature-dependent fluorescence decay of dimers. *J. Phys. Chem.* **95**, 6095–6104 (1991).

5. Paviolo, C., Clayton, A. H. A., Mcarthur, S. L. & Stoddart, P. R. Temperature measurement in the microscopic regime: A comparison between fluorescence lifetime- and intensity-based methods. *J. Microsc.* **250**, 179–188 (2013).

6. Inada, N., Fukuda, N., Hayashi, T. & Uchiyama, S. Temperature imaging using a cationic linear fluorescent polymeric thermometer and fluorescence lifetime imaging microscopy. *Nat. Protoc.* **14**, 1293–1321 (2019).

7. Hayashi, T., Fukuda, N., Uchiyama, S. & Inada, N. A cell-permeable fluorescent polymeric thermometer for intracellular temperature mapping in mammalian cell lines. *PLoS ONE* **10**, e0117677 (2015).

8. Stephens, A. D., Lu, M., Fernandez-Villegas, A. & Kaminski Schierle, G. S. Fast Purification of Recombinant Monomeric Amyloid-β from E. coli and Amyloid-β-mCherry Aggregates from Mammalian Cells. *ACS Chem. Neurosci* **11**, 3213 (2020).

9. Ovesný, M., Křížek, P., Borkovec, J., Švindrych, Z. & Hagen, G. M. ThunderSTORM: A comprehensive ImageJ plug-in for PALM and STORM data analysis and super-resolution imaging. *Bioinf.* **30**, 2389–2390 (2014).
